# Sex-dependent plasticity of adult neural tissue in response to damage

**DOI:** 10.1101/2025.05.05.652027

**Authors:** Marina Recatalà-Martinez, Manel Bosch, Pedro Gaspar, Alessandro Mineo, Santiago Rios, Irene Miguel-Aliaga, Marta Morey

## Abstract

Plasticity of intact adult neural tissue in the vicinity of neural damage serves to restore functionality of circuits. Much remains to be learned about the mechanisms regulating this process and the reported sex differences in recovery outcomes. Here, we present the fly gut and its innervation as a simplified model to address these questions. We show that gut damage caused by ingestion of toxic agents resulted in plasticity of the adult enteric neuronal network, manifested as a Reactive Oxygen Species (ROS)-dependent increase in neural tissue, which was reversible after a recovery period. Interestingly, males did not display neural plasticity, and masculinization of neurons in females suppressed the damage-dependent neural growth. Together, these findings position the fly gut as a system to investigate the cellular, molecular, and sex-specific underpinnings of neural plasticity, with implications for therapeutic advancements in neural circuit recovery.

**SUMMARY STATEMENT:** This study establishes the fly gut as a simple system to explore how adult neural tissues differ between sexes in their capacity for plasticity.

## INTRODUCTION

Neural plasticity involves two key mechanisms for recovery after injury: axonal regeneration and structural plasticity of intact neural tissue. While axonal regeneration has been widely studied (Mahar and Cavalli, 2018; Tedeschi and Bradke, 2017), modulating the plasticity of remaining healthy neural tissue offers an alternative route to restore function (Gao et al., 2022; Petersen et al., 2022). Clinical evidence shows that this form of plasticity supports functional recovery and circuit reorganization after acute damage such as stroke (Cirillo et al., 2020; Murphy and Corbett, 2009; Nudo, 2003; Sampaio-Baptista et al., 2018) or multiple sclerosis episodes (Ksiazek-Winiarek et al., 2015; Prosperini et al., 2015). Notably, recovery outcomes show sex-specific differences (Turtzo and McCullough, 2008; Yu et al., 2015), suggesting variability in plasticity capacity (Hyer et al., 2018; Kirby et al., 2024).

Despite these observations, the molecular mechanisms regulating plasticity in intact adult neural tissue remain poorly understood. This is partly due to the technical challenges of studying healthy and damaged neurons in close proximity, which make it difficult to isolate and study specific structural plasticity processes. To address this, we developed a model that focuses on how intact neurons respond to non-neural tissue damage, using peripheral innervation as an accessible system. This allows for controlled injury and analysis of plasticity in a physiologically relevant yet simplified context.

We used the digestive tract of *Drosophila melanogaster* as a model system. Like in mammals, the fly gut is structurally and functionally compartmentalized (Buchon et al., 2013; Lemaitre and Miguel-Aliaga, 2013; Marianes and Spradling, 2013). Its epithelium includes intestinal stem cells (ISCs), enteroblasts (EBs), absorptive enterocytes (ECs), and secretory enteroendocrine (EE) cells. The gut is wrapped in muscle, oxygenated by a tracheal system, and innervated by neurons located in the central nervous system, the corpora cardiaca (neurosecretory structure), and the hypocerebral ganglion (HCG, one of the stomatogastric ganglia) (Kuraishi et al., 2015; Miguel-Aliaga et al., 2018). While most neurites target visceral muscle, some extend to the underlying epithelium (Cognigni et al., 2011; Cui et al., 2024; Kenmoku et al., 2016; Petsakou et al., 2023). Unlike mammals, innervation in the fly gut is regionally restricted, making it well-suited for targeted analysis (Kuraishi et al., 2015; Miguel-Aliaga et al., 2018). Recent studies reveal that gastrointestinal neurons respond to factors such as microbiota, nutrients, aging, and reproductive state in flies and mammals (Ameku et al., 2020; Hadjieconomou et al., 2020). While functional plasticity has been observed in the posterior fly midgut during recovery (Petsakou et al., 2023), the mechanisms of structural plasticity in response to gut damage—and how they are shaped by biological sex—remain unexplored. Here, we show that intact neurons innervating the anterior midgut exhibit a female-specific, ROS-dependent, reversible structural plasticity, which is absent in male flies and suppressed in females with masculinized neurons.

## RESULTS AND DISCUSSION

### Intestinal damage induces reversible neural plasticity

In order to quantify and characterize the innervation in the anterior part of the midgut, we proceeded to select a neural driver and reporter that would specifically label neuronal processes with native GFP fluorescence. This prevented background signal from secondary antibodies and provided a high signal-to-noise ratio, which facilitated faithful neural process quantification as GFP signal. General neuronal markers, such as Elav (Chen et al., 2016; Holsopple et al., 2022; Titos et al., 2023) or n-Synaptobrevin (Weaver et al., 2020), also label EEs. However, we used the *GMR51F12-GAL4* driver line from the Janelia FlyLight collection (Jenett et al., 2012; Min et al., 2021; Olds and Xu, 2014) and a specific *nSyb.S-GAL4* driver (Weaver et al., 2020), neither of which labels EEs. Morphological landmarks in whole-mount preparations, combined with phalloidin co-staining to visualize the surrounding musculature, enabled consistent imaging of the same digestive tract region and its associated neural innervation (Fig. 1A). Using 3D reconstruction and image analysis, we obtained volume quantifications for both the digestive tube and neural tissue (see Materials and Methods). Scaling growth is a prevalent phenomenon in the peripheral nervous system, where sensory and motor neurons must adjust the size of their arborizations according to the area of their target tissues to maintain their functionality (Bentley and Toroian-Raymond, 1981; Bucher and Pflüger, 2000; Lee and Stevens, 2007; Menon et al., 2013). Measuring the volume of the digestive tube enabled us to use statistical analysis methods that accounted for and removed any potential influence of digestive tube size variation in adults (driven by treatment or genetic background) on neural tissue quantification comparisons and distinguish scaling growth from neural plasticity tissue growth (see Materials and Methods).

**Fig. 1.**
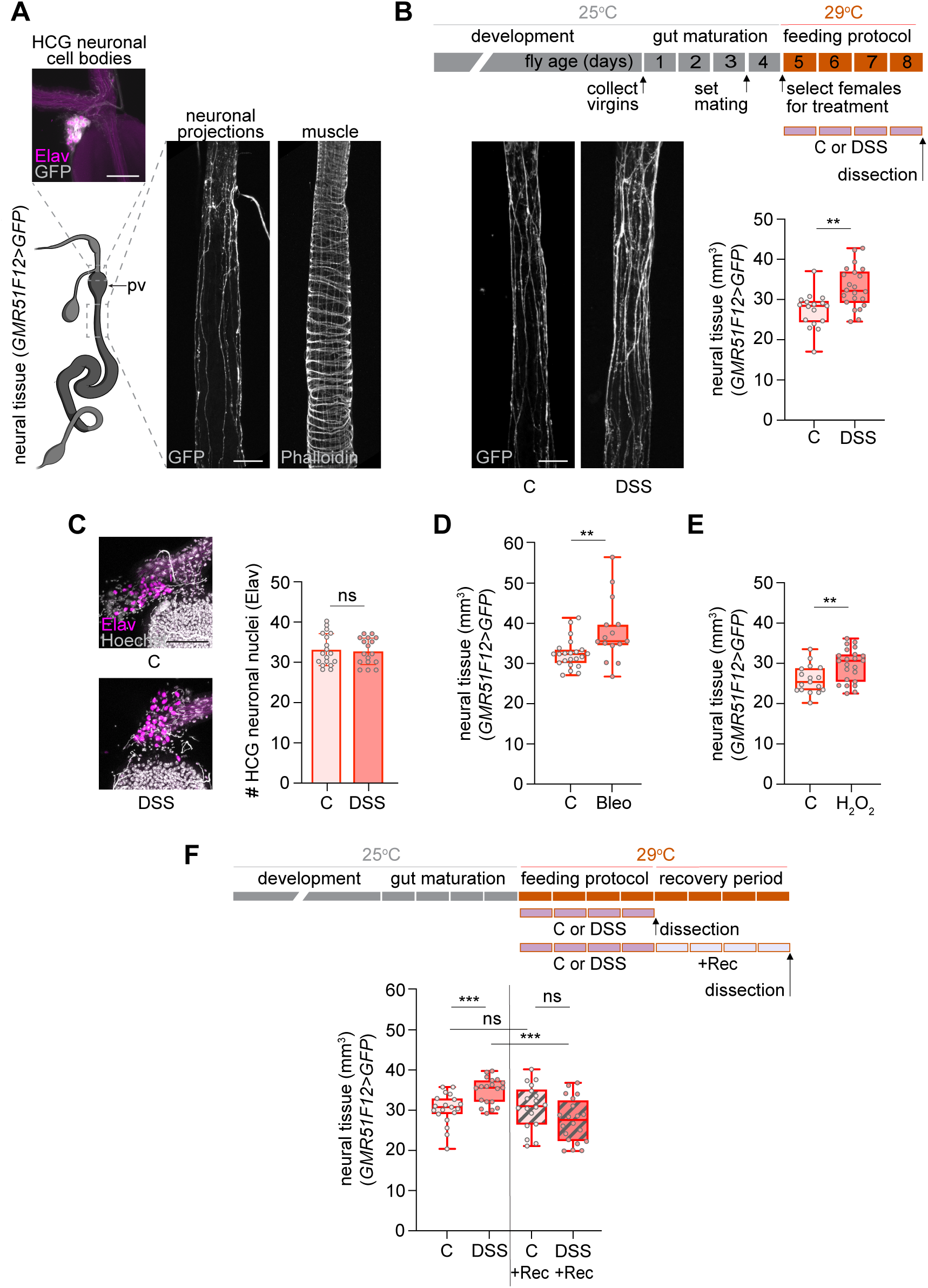
Neural plasticity induced by gut damage resolves after recovery. A) Drawing of the adult gut with two areas of interest highlighted and corresponding confocal Z-projections. The square box region corresponds to the area above the proventriculus (pv) and the image shows neuronal nuclei (magenta, anti-Elav) and neuronal membrane (grey, GFP) of the HCG. Autofluorescence of the peritrophic membrane lining the gut in this region is observed. The rectangular box region corresponds to the area of the anterior midgut imaged in subsequent analyses and the images show neuronal processes (grey, GFP) and muscle fibers (grey, Phalloidin). B) Detailed schematic outlining the experiment in the panel, with rectangles representing 1 day each and reflecting the age of the flies (brown) and the duration of the feeding protocol (purple) or any other additional period after that (light purple, like in panel F). This type of schematic is used throughout the figures to describe the experiment carried out. Confocal Z-projections of neuronal processes innervating the anterior midgut of control (C) and DSS-fed (DSS) females and quantification of neural tissue. Red outline in the box plots or bar graphs indicates female. C) Confocal Z-projections of the HCG of C and DSS-fed females stained with Hoechst (grey, which also accumulates in some tracheal branches) and showing neuronal nuclei (magenta, anti-Elav). Quantification of neuronal nuclei in the HCG of C and DSS-fed females. D) Quantification of neural tissue in C and Bleomycin (Bleo)-fed females. E) Quantification of neural tissue in C and H2O2-fed females. F) Schematic of the feeding protocol and recovery period. Quantification of neural tissue after the feeding protocol and after the recovery period (Rec). Scale bar 50 μm. Statistical significance is indicated as follows: *\* p < 0.05, ** p < 0.01*, *\*\*\* p < 0.001,* ns = not significant. Gut schematic from *BioRender.com*.

To damage the gut, we fed flies with DSS. DSS is a polymer that induces colitis in mammals (Yang and Merlin, 2024) and has been used in *Drosophila* to study the cellular mechanisms and molecular pathways that regulate ISC proliferation during regeneration (Jiang et al., 2016; Tian et al., 2022). When flies ingest DSS, it crosses the epithelium, expanding the basement membrane sheet and altering muscle morphology (Howard et al., 2019). The resulting change in biomechanical forces in the niche is thought to activate ISC division (Howard et al., 2019). DSS feeding also results in trachea sprouting (Perochon et al., 2021), which is necessary for ISC proliferation (Medina et al., 2025; Perochon et al., 2021; Tamamouna et al., 2021). Interestingly, basement membrane and muscle defects, as well as trachea remodeling, are reversible after a recovery period (Howard et al., 2019; Perochon et al., 2021; Tamamouna et al., 2021). We wondered whether adult neurons were plastic and could also respond to gut damage.

To analyze the effect of gut damage on neural plasticity, we fed adult mated females DSS for 4 days and compared them to controls fed with the carrier (sucrose) (Fig. S1A-D). We confirmed the effectiveness of DSS by replicating previously published observations, including a reduction in lifespan (Amcheslavsky et al., 2009) (data not shown), and an increase in the proliferative response of ISCs in the midgut (Amcheslavsky et al., 2009) (see later in Fig. 2A). Over 90% of the flies that survived the feeding protocol tested negative for the SMURF assay (Rera et al., 2012), indicating that in the majority of animals (93,3%±SD4.5) that were dissected gut permeability and integrity were not compromised (Fig. S1E,F). Thus, during DSS feeding, neurons were largely shielded from direct exposure to gut contents that could potentially cause them damage. When we dissected the DSS-fed females and compared them to control flies, we observed a significant increase in neural tissue (Fig. 1B). Virgin flies also showed a DSS-dependent increase in neural tissue, comparable in magnitude to that observed in mated females (Fig. S1H,I). To rule out neurogenesis as the source of the new neural tissue, we quantified the number of neural nuclei in the HCG, the primary source of neurons innervating the anterior midgut (Fig. 1A). No differences were observed between control and DSS-fed animals (Fig. 1C), suggesting that the increase in neural tissue was due to neural growth. This neural growth was not exclusive to DSS damage since we also observed it when feeding the flies Bleomycin and H2O2 (Fig. 1D,E, Fig. S2A,B). In contrast, *Pseudomonas entomophila* infection led to a marked gut enlargement in our region of interest, with neural tissue scaling accordingly. Because this expansion likely approached the limit of neural growth, it prevented detection of any plasticity growth beyond this scaling response (Fig. S2C). Overall, several forms of chemically induced damage paradigms robustly trigger neural growth.

**Fig. 2.**
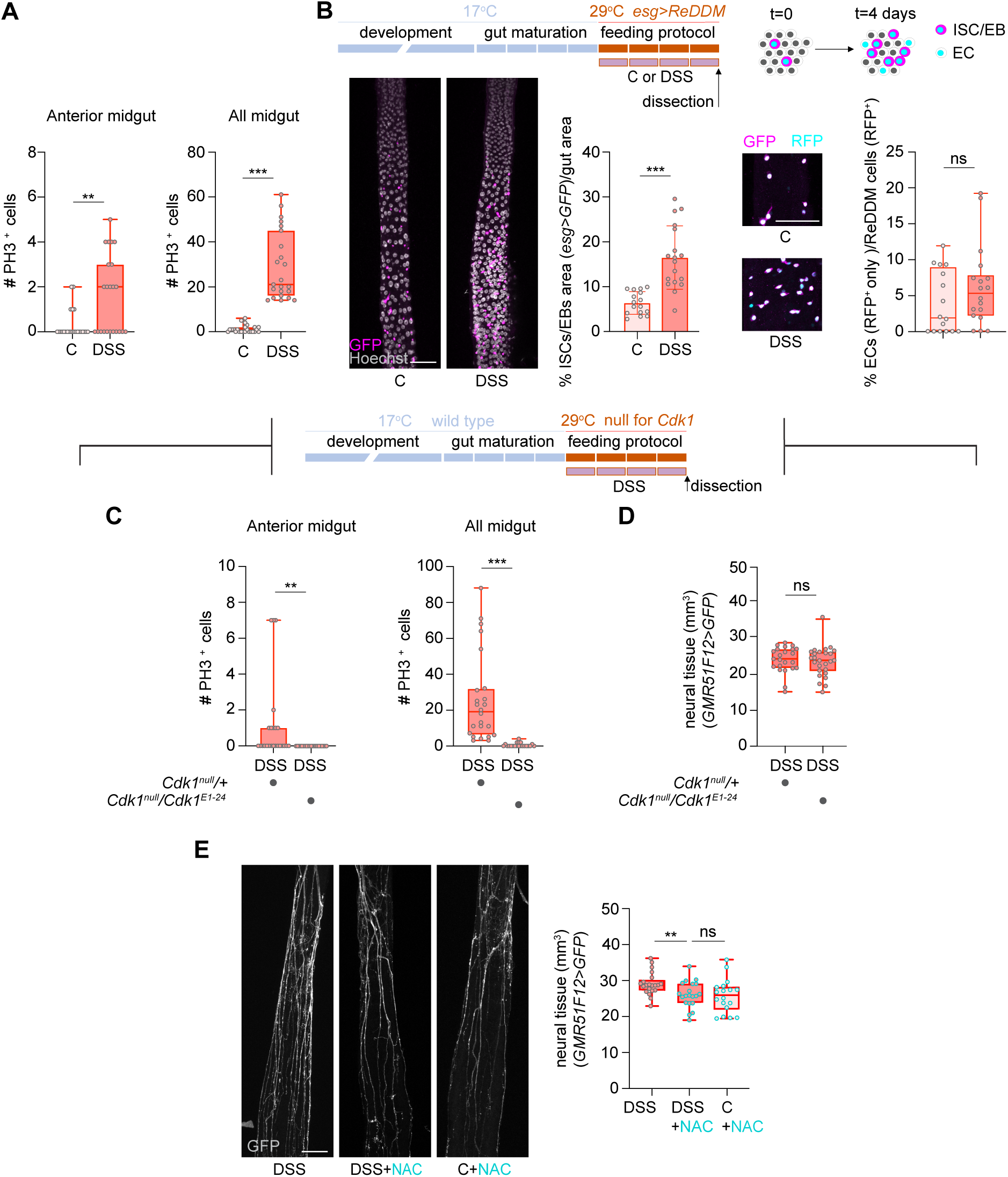
Neural plasticity does not depend on ISC proliferation but on gut-derived ROS. A) Quantification of PH3/Hoechst positive cells in the anterior and the whole midgut in C and DSS-fed females. B) Schematic showing activation of ReDDM lineage tracing during the feeding protocol and cell types distinguished in the lineage. Confocal Z-projections of the anterior midgut: stained with Hoechst (grey) and showing the cell membrane of ISCs/EBs (magenta, anti-GFP); and showing differentiated ECs (cyan, RFP^+^ only) in C and DSS-fed females. Quantification of the percentage of the ISC/EB area (GFP signal) from the total area of the anterior mid gut and of the differentiated ECs (RFP^+^ only) from ReDDM cells (RFP^+^) in C and DSS-fed females. Although not statistically significant, the increased percentage of ECs in the anterior midgut in DSS-fed animals was a consistent trend in biological replicates of this experiment. C-D) Schematic showing temperature-dependent knock-out of Cdk1 function during the feeding protocol. C) Quantification of PH3/Hoechst positive cells in the anterior midgut and the whole midgut in wild type (heterozygous) and *Cdk1* null females after DSS feeding. D) Quantification of neural tissue in wild-type and *Cdk1* null females after DSS feeding. E) Confocal Z-projections of neuronal processes in the anterior midgut of DSS, DSS + NAC and C +NAC -fed females and quantification of neural tissue. Scale bar 50 μm. Statistical significance is indicated as follows: *\* p < 0.05, ** p < 0.01*, *\*\*\* p < 0.001,* ns = not significant.

Tracheal terminal cells (TTCs) sprout in response to DSS, Bleomycin, H2O2, and infectious pathogens (Medina et al., 2025; Perochon et al., 2021; Tamamouna et al., 2021), so we examined their behavior in the anterior midgut. This region showed sparse tracheation, which remained unchanged after DSS feeding, indicating that TTCs in this area were unresponsive to gut damage (Fig. S3A, B). Moreover, TTC morphology did not match the extent of neural growth, suggesting that they do not serve as a scaffold for neural tissue growth in this area.

We next explored the stability of this neural plasticity and conducted recovery experiments (Fig. 1F). While DSS feeding increased the neural tissue (C versus DSS), a 4-day recovery period (DSS + Rec) reverted the amount of neural tissue to levels as those of control flies (C + Rec). Consistent with this observation, a clear reversal of neural growth was observed when comparing flies dissected after the DSS treatment (DSS) and those dissected after the recovery phase (DSS + Rec). No differences were observed between control flies (C versus C + Rec).

In summary, these experiments indicate that the intact neural tissue innervating the digestive tube is plastic and responds to damage with growth. This neural growth is reversible after a recovery period, as the gut returns to homeostasis after injury. Importantly, these observations were validated with two independent neural drivers (*GMR51F12-GAL4*, Fig. 1 and *nSyb.S-GAL4*, Fig. S3C).

### ISC proliferation does not mediate neural plasticity

To determine whether ISC proliferation drove DSS-induced neural growth, we first confirmed that DSS feeding increased ISC division in the anterior midgut, as shown by elevated phospho-histone H3 (PH3) signal (Fig. 2A) and lineage tracing using ReDDM (Antonello et al., 2015) (Fig. 2B) (see Materials and Methods). To test causality, we blocked ISC cell division during DSS feeding using a combination of a *Cdk1* null allele (*Cdk1^null^*) and a temperature-sensitive allele (*Cdk^1E1–24^*) (Rodríguez et al., 2024) (Fig. 2C). Neural growth remained unaffected by the proliferation block (Fig. 2D), indicating that ISC division is not required for DSS-induced neural plasticity.

### Gut-derived ROS stimulates neural plasticity

DSS feeding results in epithelial damage, causing cell death and an increase in ROS levels (Wu et al., 2012)(Fig. S4A). To test the contribution of ROS levels to neural growth, we co-fed flies with DSS and the ROS inhibitor N-acetyl cysteine (NAC). Neural growth was reduced in DSS + NAC flies and comparable to control + NAC levels (Fig. 2E), indicating that DSS-dependent ROS promotes neural plasticity. This finding aligns with evidence that ROS supports structural plasticity (Oswald et al., 2018).

It has been recently shown that ROS from midgut epithelial cells promotes the secretion of diuretic hormone 31 (Dh31) from anterior EE cells and that Dh31 binds Dh31 receptor (Dh31-R) to mediate tracheal remodeling (Medina et al., 2025). To test whether ROS-induced Dh31 signaling mediates the increase in neural tissue observed after DSS feeding, we verified that Dh31-R was expressed in neurons in the HCG (Medina et al., 2025)(Fig. S4C). We analyzed *Dh31* mutants and flies with neuronal knockdown of its receptor, *Dh31-R* (Fig. S4D-G). Baseline innervation was unaffected in either condition (Fig. S4D,F). DSS feeding induced a comparable increase in neural tissue in *Dh31* homozygous mutants and heterozygous controls (Fig. S4E), as well as in *Dh31R* knockdown flies (Fig. S4G). These results indicate that the DSS-dependent, ROS-induced neural plasticity occurs independently of the Dh31/Dh31-R pathway.

### Reversal of neural growth as the gut returns to homeostasis is not caspase dependent

We were curious about the mechanisms controlling neural growth reversal during recovery. Axon pruning can take place either through axon retraction or degeneration (Luo and O’Leary, 2005). During developmental remodeling, dendritic arborizing (da) sensory neurons ddaC degenerate (Williams and Truman, 2005) using local caspase activation to direct dendrite engulfment (Williams et al., 2006). To assess whether caspases mediated neural growth reversal, we inhibited apoptosis in neurons by overexpressing Diap1. Reversal still occurred (Fig. S5), suggesting that this process may result from axon retraction rather than caspase-dependent mechanisms.

### Neural plasticity is exclusive to females and is actively suppressed when female neurons are masculinized

Interestingly, males did not exhibit neural growth following DSS treatment (Fig. 3A) despite a DSS-dependent increase in ROS (Fig. S4B) and reduced viability (Amcheslavsky et al., 2009). To investigate whether this neural growth was exclusive to female neurons, we aimed to create female individuals with masculinized neurons. Most sex differences in *Drosophila* arise from the female-specific expression of the *Sex lethal* gene (*Sxl*), which codes for an RNA-binding protein (Bell et al., 1988). Sxl induces the alternative splicing of the *transformer* (*tra*) gene, generating the female fate-determining TraF protein only in females (Belote et al., 1989; Boggs et al., 1987; Inoue et al., 1990; Sosnowski et al., 1989). We first confirmed that neurons in the HCG expressed the female-specific alternative splicing variant of *tra* while males didn’t (Fig. 3B,C) (Hérault et al., 2024). Subsequently, we proceeded to masculinize neurons in females by means of *tra* downregulation with RNAi during development (Fig. 3D). We confirmed that the presence of the *UAS-Dcr2* transgene did not affect the response of wild-type females to DSS, and thus, the expected increase in neural tissue was observed. Interestingly, DSS feeding did not have any effect on masculinized females, and the amount of neural tissue was not significantly different between control and DSS-fed flies. In addition, DSS-fed masculinized females displayed significantly less neural tissue than DSS-fed control flies, aligning with the absence of neural growth in masculinized females.

**Fig. 3.**
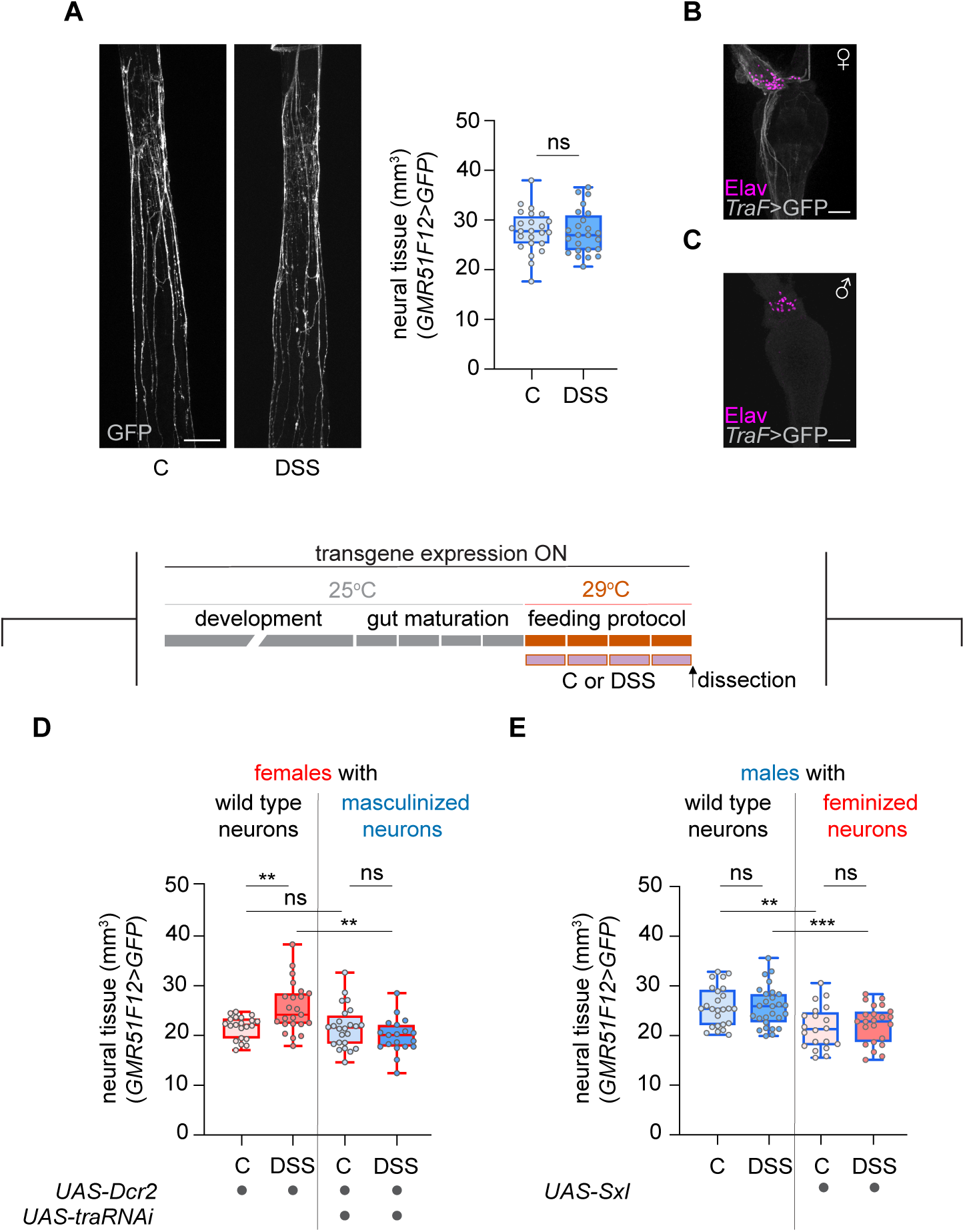
Neural plasticity is not observed in males and masculinization of female neurons abolishes neural growth. A) Confocal Z-projections of neuronal processes in the anterior midgut of C and DSS-fed males, and quantification of neural tissue. Blue outline in box plots indicates male. B-C) Confocal Z-projections at the level of the HCG showing female-specific *tra* alternative splicing in neurons (grey, GFP; magenta, Elav) in females (B) and absence in males (C). D-E) Schematic showing transgene expression in neurons in relation to the feeding protocol. D) Quantification and comparison of neural tissue in females and females with masculinized neurons both in C versus DSS-fed conditions. E) Quantification and comparison of neural tissue in males and males with feminized neurons both in C versus DSS-fed conditions. Scale bar 50 μm. Statistical significance is indicated as follows: *\* p < 0.05, ** p < 0.01*, *\*\*\* p < 0.001,* ns = not significant

We also carried out the reverse experiment and attempted to confer plasticity to male neurons by feminizing them through developmental *Sxl* overexpression (Fig. 3E) (Bell et al., 1991). In this case, the DSS treatment had no effect on feminized males, and we did not observe the neural growth we expected to see for a successfully feminized male. Moreover, both control and DSS-fed feminized males showed less amount of neural tissue than wild-type males. These results indicate that neuronal Sxl overexpression in males is not sufficient to induce damage-responsive plasticity and may even impair neural tissue growth in our context.

These findings reveal a previously unrecognized sex-specific mechanism of neural plasticity in the fly gut, where female neurons exhibit a unique capacity for damage-induced growth that is actively suppressed by masculinization. In addition, the lack of neural growth in males following DSS-induced gut damage suggests that neural plasticity is limited in males compared to females.

### Female-specific neural plasticity impacts gut physiology and viability

To address the functional impact of neural plasticity, we carried out defecation assays both after DSS treatment and after recovery treatment. We observed that DSS feeding caused a robust increase in defecation (Fig. 4A,B) (see Material and Methods). After a 4-day recovery period on sucrose (Fig. 4A), defecation in flies previously exposed to DSS (DSS + Rec) remained significantly higher than in control flies (C + Rec), although neural tissue increase reverts in these conditions (Fig. 1F). When the recovery period was performed on standard food (Fig. 4B), defecation in DSS-treated flies (DSS + Rec) was not significantly different from controls (C + Rec). Although the difference under this condition did not reach statistical significance, the values still showed a mild upward trend in DSS-recovered flies, which is consistent with the expectation that physiology may not be fully restored but approaches near-normal levels. These observations suggest that standard food promotes recovery of the digestive tube, and that normalization of defecation requires coordinated restoration of both neural and gut homeostasis.

**Fig. 4.**
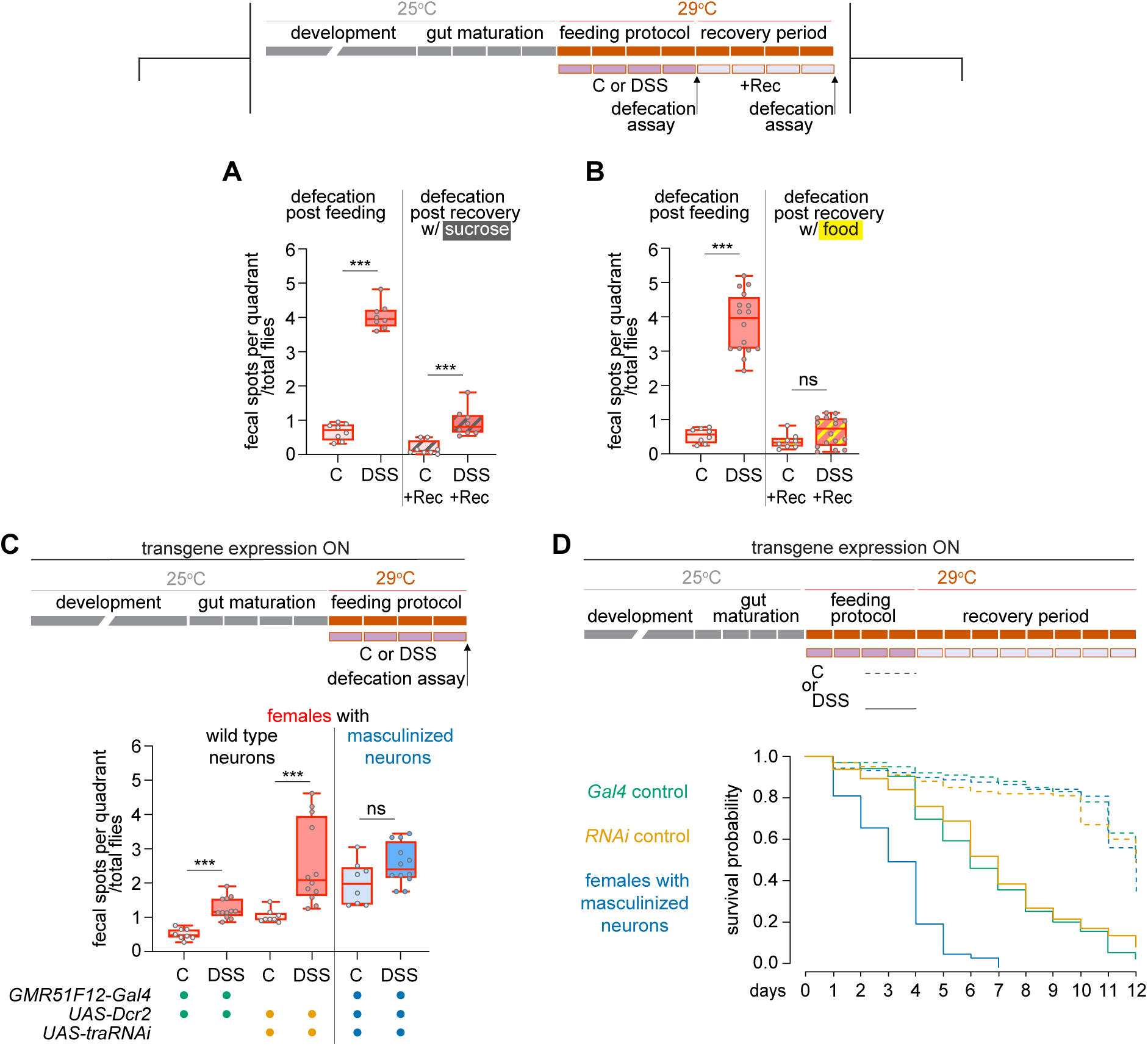
Neural plasticity in females is required for DSS-induced defecation increase and protects against DSS-induced mortality. A, B) Schematic outlining the timing of defecation assays. Quantification of defecation in a cohort of flies after C or DSS feeding, and after a recovery period: A) on 5% sucrose, or B) on regular food. C) Schematic showing transgene expression and timing of defecation assay. Quantification of defecation after C or DSS feeding in females with wild-type or masculinized neurons. For females with wild-type neurons, controls for the *Gal4/dcr2* and *RNAi/dcr2* transgenes are shown. The three genotypes are color-coded: *Gal4* control in green, *RNAi* control in yellow, and females with masculinized neurons in blue. D) Schematic showing that lifespan analysis covers the 4 days of the feeding assay plus 8 additional of recovery where flies were fed on sucrose. Survival curves of females after C or DSS feeding with wild-type or masculinized neurons, color-coded as in panel C. When DSS-fed, females with masculinized neurons have a significantly reduced life span compared to females with wild-type neurons (p value=<0,001 for both comparison to *Gal4/dcr2* control and to *RNAi/dcr2* control). For additional comparisons, see Table S2. Statistical significance is indicated as follows: *\* p < 0.05, ** p < 0.01*, *\*\*\* p < 0.001,* ns = not significant.

However, whether neural plasticity contributes to the increased defecation and its potential protective role remained unclear. To address this, we masculinized female neurons to block their DSS-induced plasticity. This manipulation markedly suppressed the defecation increase after DSS feeding (Fig. 4C), indicating that neural plasticity is required for this physiological response. Independently, lifespan analysis showed that eliminating neural plasticity (i.e. masculinizating neurons) further reduced the survival of DSS-fed female flies (Fig. 4D), suggesting that the plastic response also provides a protective benefit.

Together, these results reveal two different effects of neural plasticity under DSS challenge, even though how the increase in neural tissue contributes to these behaviors and whether they are connected remains unknown. Nonetheless, they highlight the importance of neural plasticity in shaping the physiological response to gut damage.

### The *Drosophila* gut and its innervation as a model to investigate neural plasticity and the role of sex in its regulation

This system enables the study of both structural growth and retraction phases of adult intact neural tissue in response to damage, capturing the dynamic nature of plasticity. In addition, we uncovered a clear sexual dimorphism: neural growth is absent in males and suppressed in masculinized female neurons. However, feminizing male neurons was not sufficient to induce plasticity.

Two complementary directions emerge for understanding this dimorphism. One line of inquiry concerns sex-specific gene expression that may underlie the differential plasticity. Sex differences in genes regulating ROS production and/or ROS-responsive signalling pathways could drive the female-specific neural growth observed in response to gut damage. For the latter, candidate pathways include TOR and InR/PI3K/Akt, which regulate neural regrowth during developmental remodeling in flies (Sanal et al., 2023; Yaniv and Schuldiner, 2016) and mammalian regeneration (Lund-Ricard et al., 2020). Other molecules downstream of ROS are Umpaired (Upd) cytokines, such as Upd3 (Santabárbara-Ruiz et al., 2015), which could link tissue damage to neural plasticity (see below).

The other direction focuses on the influence of surrounding tissues on neural plasticity. The failure of feminized male neurons to elicit growth in response to damage implies that neuronal identity alone is not enough—sex-specific cues from surrounding tissues such as the gut epithelium, visceral muscle, tracheal cells or Malpighian tubules may be necessary to support or trigger neural growth. Indeed, Upd3 can be secreted by damaged gut epithelial cells, visceral muscle or Malpighian tubules, leading directly or indirectly to ISC proliferation (Buchon et al., 2010, 2009a, 2009b; Jiang et al., 2009; Lin et al., 2010; Liu et al., 2024; Medina et al., 2022; Osman et al., 2012; Tian et al., 2015). Recent studies show that neurons can respond directly to glia-derived Upd3, which modulates neuronal structure (Sodders et al., 2025), and indirectly to gut-derived Upd2/3 via glial intermediates, which influences neuronal function (Malita et al., 2025). Together, these initial insights raise the possibility that the neural plasticity observed after DSS feeding may be shaped by Unpaired ligands present in the gut microenvironment. Similarly, reversal of neural growth during recovery may involve repulsive cues such as Ephrins, Slits, or Semaphorins, potentially transiently expressed in tissues of the digestive tube environment in females, thereby mediating axon retraction in a manner reminiscent of classic axon-guidance mechanisms (Luo and O’Leary, 2005).

Building on evidence that sex influences organ formation and physiology (Mank and Rideout, 2021), the results presented here establish the *Drosophila* gut and its innervation as a model for elucidating how sex shapes neural plasticity of intact neural tissue in response to damage. Understanding these mechanisms could guide the development of sex-specific therapeutic strategies for neural repair.

## MATERIALS AND METHODS

### Fly stocks

The following fly lines were used: *GMR51F12-GAL4* (BDSC:58685), *nSyb.S-GAL4* (BDSC: 51635), *UAS-Flybow.1.1B* (used as 10xUAS-CD8::GFP, BDSC: 56802, 56803), *DSRF.Term-GAL4* (BDSC: 25753), *esg-GAL4* (a gift from M. Milán, unknown insertion), *UAS-mCD8::GFP* (BDSC:5130), *UAS-H2B::RFP* (a gift from J. Morante), *tubP-GAL80^ts^* (BDSC: 7017), *Cdk1^null^* (BDSC: 6643), *Cdk^1E1–24^* (BDSC: 6641), *Dh31^attP^*(BDSC: 84490), *UAS-Dh31-R RNAi TRIPJF01945* (UAS-Dh31-R RNAi, BDSC: 25925), *Dh31-R^2A-Gal4^* (BDSC: 84626), *UAS-Diap1* (BDSC: 15310), *traF-GAL4* (a gift from B. Hudry), *UAS-Dcr-2* (BDSC: 24650), *UAS-tra RNAi TRiPJF03132* (*UAS-tra RNAi*, BDSC: 28512), *UASp-Sxl.alt5-C8* (*UAS-Sxl*, BDSC: 58484). The list of complete genotypes is available in Supplementary Information.

### Feeding protocols

For feeding assays, virgin females were collected and aged on standard media at 25°C for 3-4 days before undergoing mating for 24 h. The same procedure was carried out when experiments were performed with males. Flies were recovered from the mating and transferred in groups of 10 to an empty vial containing 5 pieces of 2cm × 2cm of bench paper soaked with 500ml of the desired feeding solution. Flies were transferred to a new vial with fresh feeding paper every day for the specified number of days prior to dissection. The feeding protocol was always done at 29°C, all feeding treatments were administered in 5% sucrose (S7903, Sigma Aldrich), and flies fed with sucrose alone served as the control (C) condition unless otherwise stated. Flies were fed with DSS (42867, Sigma Aldrich) at 6% w/v for 4 days. For the SMURF assay, the last control and DSS feeding solutions prior to the day of dissection were laced with Brilliant Blue FCF (80717, Supelco) at 2.5% w/v. Bleomycin (BL7216, Sigma Aldrich) was administered at 50 µg ml^-1^ for 3 days, H2O2 (1072100250, Supelco) at 1% for 4 days, and *Pseudomonas entomophila* (*Pe*) at an OD600=25 for 16 h. For NAC (A7250, Sigma Aldrich) co-treatment, flies were fed with 1,2 mM NAC in 5% sucrose (Control + NAC) or 6% DSS + 1,2 mM NAC in 5% sucrose (DSS + NAC) for 4 days. For recovery (Rec) experiments, after the feeding protocol, C and DSS-fed flies were each transferred to vials with 5% sucrose for 4 days (C + Rec ; DSS + Rec). To carry out comparisons within the same cohort of flies, a subset of flies was dissected after the feeding protocol, and the rest after the recovery period.

### Temperature-sensitive assays

To visualize the amount of proliferation taking place exclusively during the four days of the feeding protocol, we set ReDDM-based crosses (Antonello et al., 2015). This lineage tracing approach uses the Gal80 temperature-sensitive allele (*Gal80^ts^*), which inhibits GAL4 transcription at permissive temperature (19°C) and allows it at the restrictive one (29°C) (McGuire et al., 2003), and *esg-GAL4* to label ISCs and EBs. Crosses were grown and flies were kept at 17°C to ensure Gal80^ts^ repression until adult mated female flies were shifted to 29°C to inactivate Gal80 and carry out the feeding protocol. To block ISC proliferation during the feeding protocol, *Cdk1^null^/ Cdk^1E1–24^* flies were grown, aged, and mated at 17°C, Cdk1 function was abrogated once the flies were transferred to 29°C (Rodriguez et al., 2024) during the DSS feeding protocol.

### Dissection and immunostaining

To obtain the gut, flies were dissected in PBS. Adult animals had their heads cut off with scissors by the neck, their abdomens were pulled apart from the thorax, and opened to extract the gut. With this procedure, the digestive tube was severed well above the crop duct. Once the fat and ovaries were removed, the gut was handled by its posterior end and transferred to PBS in a well drawn onto a poly-L-lysine-coated slide (P1524, Merck) using hydrophobic silicone (Flowsil, Intek Adhesives). In cases where the anterior part of the midgut had to be repositioned, we manipulated the gut from the crop to avoid touching the digestive tube. Guts were fixed at room temperature for 30 min in 4% PFA in PBS. Washes were done with PBS-T (PBS, 0.25% Triton X-100). To label the muscle or nuclei, guts were incubated at room temperature for 30 min with Alexa Fluor^TM^ 635 Phalloidin (1/400, A34054, Invitrogen) or Hoechst 33258 (1/1000, H3569, Invitrogen), respectively, in PBS-T. A final wash with PBS was done before mounting. When antibodies were used, primaries were incubated overnight at 4°C, and secondaries were incubated at room temperature for 2–3 h. The following antibodies were used: rat-anti-Elav (1/100, 7E8A10, DSHB), mouse-anti-Discs large (1/50, 4F3, DSHB), rabbit-anti-PH3 (1/1000, 06-570, EMD Milipore), rabbit-anti-DsRed (1/200, 632496, Clontech), chicken-anti-GFP (1/800, ab13970, Abcam), Alexa Fluor^TM^ 568 goat-anti-rat, Alexa Fluor^TM^ 488 goat-anti-mouse, Alexa Fluor^TM^ 488 goat-anti-rabbit, Alexa Fluor^TM^ 568 goat-anti-rabbit, Alexa Fluor^TM^ 488 goat-anti-chicken (1/500, A11077, A11001, A11008, A11011, A11039, Life Technologies). Samples were mounted in VECTASHIELD® PLUS Antifade Mounting (H-1900-10, Vector Laboratories).

### ROS detection

C and DSS-fed guts were dissected in Schneider’s medium (50146, Sigma Aldrich) and placed on poly-L-lysine-coated slides. Tissues were incubated in CM-H2DCFDA (C6827, Invitrogen) at 2 µM in Schneider’s medium for 30 min in the dark, then washed three times in Schneider’s medium before imaging. Dissection time was limited to 15 min for each experimental group, and imaging was performed within 45 min of dissection.

### Defecation assay

Flies under analysis were starved for 1 h in empty vials. Subsequently, 20–25 flies were introduced into fly cages positioned over a 60-mm Petri dish containing a central 5 × 5 mm piece of grape-juice agar with 2.5% (w/v) Brilliant Blue FCF (Supelco, 80717). Flies were kept at 29 °C for 12 h in constant darkness before quantification of the fecal spots.

### Lifespan experiments

Lifespan analysis of females aged and mated started on the 1^st^ day of the feeding protocol and after that continued for 8 additional days on 5% sucrose. Flies were placed in groups of maximum 10 per vial, and transferred to new feeding vials every day.

### Microscopy

Samples were imaged using a Zeiss LSM 880 confocal microscope (Zeiss, Germany) equipped with a 25X 0.8 glycerol immersion media objective and an Argon and a HeNe633 lasers. For neural network analysis, to consistently image the same region in each gut, we used the end of the proventriculus as a positional landmark and captured the gut section that fit within the visual field of the 25X objective at 0.8 magnification, which in our setup generated images of 566,79 µm². As quality control for dissection, only guts with an intact HCG were considered since this ganglion contains the cell bodies of many of the neurons innervating the anterior midgut. Stacks of images were acquired with a pixel size of 0.55x0.55x1um (xyz) and a pixel dwell time of 2.05us.

### Image processing and quantification based on images

Images were processed with Fiji (version ImageJ 1.53c, (Schindelin et al., 2012) and figures assembled using Adobe Illustrator (Adobe, San Jose, CA).

*Quantification of neural tissue and gut parameters:* The channel with the neuron staining was filtered in 3D (Ollion et al., 2013) first with the maximum filter and then with the minimum one, with a radius of 3.3um in x and y and 2um in z, in both filters. Then the filtered stack was processed using the Tubeness plugin (sigma = 0.66) (Sato et al., 1998). Finally, images were binarized using the Bernsen’s AutoLocal threshold method (radius = 2) (Bernsen, 1986). The resulting binary stack was analyzed first with the 3D Object Counter function to obtain the total neuronal network volume (mm^3^).

The channel with the gut staining was processed by background subtraction and then thresholder using the Li’s method (Li and Lee, 1993). The resulting binary stack was analyzed with the 3D Object Counter function to obtain the volume of the gut.

*Other quantifications on images:* The number of Elav/Hoechst positive nuclei in the HCG was counted from the confocal stack using the 3D Object Counter plugin. The number of TTC nuclei was counted by hand from the confocal stack maximum projection. TTCs volume was obtained using the same procedure as for neural tissue quantification. The number of PH3/ Hoechst positive nuclei in the midgut was counted directly under the confocal microscope. For quantification of cumulative proliferation using ReDDM, xyz stacks of half the gut were taken and z-projected. The Hoechst signal was manually outlined to measure the gut area. To quantify the area of GFP signal from mCD8::GFP-labeled cells (representing ISCs and EBs resulting from ISCs divisions) a Gaussian Blur filter (radius = 2.0) was applied. The image was then thresholded using the "MaxEntropy" algorithm (Kapur et al., 1985), and finally, the GFP-positive regions larger than 10μm^2^ were quantified. The GFP area was represented as a percentage of the total area. To quantify the number of differentiated ECs, the total number of ReDDM cells was quantified manually using the H2B::RFP nuclear signal, the presence or absence of cytoplasmic mCD8::GFP was tracked to calculate the percentage of differentiated ECs. For ROS signal analysis, the same region used for neural network evaluation was imaged. Z-projections combining 15 optical sections were generated using maximum intensity to capture the epithelial signal. For each gut image, mean pixel intensity was quantified in two distinct areas to account for variability across the imaged gut region.

### Other quantifications

*SMURF assay*: on the 4^th^ day of the DSS feeding protocol, we assessed the proportion of surviving flies that were positive for the SMURF phenotype. Ten tubes with 10 flies each were used in each of three biologically independent replicates.

*Defecation assay:* After incubation, the Petri dish was removed, and the four quadrants of the dish were photographed separately under a stereomicroscope. Blue fecal spots in each quadrant were manually counted using ImageJ and normalized by the total number of flies in the cage. Thus, the value obtained represented fecal spots per quadrant/total flies in the cage. Each quadrant was treated as an independent technical replicate. This approach increased the number of measurements per biological replicate, thereby improving the statistical power of the analysis while ensuring complete and accurate counting of all fecal deposits. Flies from the same cohort were used in experiments where defecation assays were carried out after DSS treatment and after recovery treatment.

*Lifespan experiments:* dead flies were counted every 24 h before being transferred to a new feeding vial.

### Statistics

Data analysis was carried out using Prism 6 (GraphPad Software Inc., San Diego, CA) and Statgraphics Centurion version 18 (Statgraphics Technologies Inc., The Plains, VA). Table S1 provides statistical details (sample size for each condition, definition of values, statistical test, comparisons and p values) for each main and supplementary figure. All tests are two-tailed. For data shown in box plots, the median is given between the first and third quartiles (ends of the box). Whiskers represent the maximum and minimum values of the data. Bar graphs display the mean, with error bars representing the standard deviation.

*For the amount of neural tissue and TTCs tissue:* To analyze the effect of the treatment on neural tissue volume and TTCs tissue volume, analysis of covariance (ANCOVA) was performed. The dependent variable was neural tissue volume or TTCs tissue volume, the fixed factor was the treatment, and the covariate was gut volume (Fig. 1B,D-F; Fig. 2D,E; Fig. 3A,D,E; Fig. S1A-D,G-H, ; Fig. S2A-C; Fig. S3B-C; Fig. S4D-G; Fig. S5).

*For other quantifications:* For pairwise comparisons of data sets following a normal distribution (Fig. 1C, Fig. S4A,B), the unpaired t-test was performed. For pairwise comparisons of data sets not following a normal distribution, we used the non-parametric Mann-Whitney test (Fig. 2A-C; Fig. 4A-C; Fig. S3A).

*For lifespan experiments*: survival distributions were compared using the non-parametric log-rank test (Kaplan and Meier, 1958), which accounts for censored data. To correct for multiple comparisons, the Holm–Bonferroni method was applied to control the family-wise error rate (Holm, 1979).

## ACKNOWLEDGEMENTS

We thank M. Corominas, C. Estella, A. Gontijo, C. González, A. González-Reyes, B. Hudry, M. Llimargas, M. Milán, J. Morante, F. Serras, M. Silies, H. Stocker, BDSC, and DSHB for reagents and members of the Corominas’ and Serras’ laboratories for insightful discussions. We are grateful to: Q. Zhu, P. Longas, and O. Ramirez for their assistance with early pilot experiments that helped shape this research; PG. Camara, P. Nicodemus, V. Pisarenco, and J. Rozas for help in exploring the use of a metric-geometry framework (CAJAL) to describe morphology changes in the anterior midgut neural network; J. Vila for help with *Pe* feeding experiments, and C. Herrera for informatic support. We are indebted to our colleagues F. Cebrià, M. Corominas, F. Serras, M. Valls, T. Adell, C. González-Estevez, L. di Croce, L. Morey, B. Payer, and the CCiTUB technological platform for their invaluable support and encouragement during this project.

## COMPETING INTERESTS

The authors declare no competing interests.

## AUTHOR CONTRIBUTIONS

Conceptualization: M.M.; Methodology: M.R-M., M.B., P.G., A.M., S.R., I.M-A., M.M.; Investigation: M.R-M., M.M.; Formal Analysis: M.R-M., P.G., A.M., S.R., I.M-A., M.M.; Validation: M.R-M., M.M.; Visualization: M.R-M., M.M.; Data Curation: M.R-M., M.M.; Resources: P.G., A.M., I.M-A.; Supervision: M.M.; Project Administration: M.M.; Funding Acquisition: M.M.; Writing – Original Draft: M.M. with contributions from all the authors; Writing- review and editing: M.M. AI-assisted tools (ChatGPT, GTP-5) were used for language editing during manuscript preparation. All content generated by the AI was reviewed and edited by the authors, who take full responsibility for the final text.

## FUNDING

This work was funded by the Spanish Ministerio de Ciencia e Innovación (grant PID2019-107340GB-I00) and Agència de Gestió d’Ajuts Universitaris i de Recerca of the Generalitat de Catalunya (2021SGR00293) to M.M. Work in I.M-A. was funded by an ERC Advanced Grant (ERCAdG 787470 “IntraGutSex”), a UKRI Horizon Europe guarantee grant (EPSRC-ERC EP/Y03298/1), MRC intramural funding and the Francis Crick Institute, which receives its core funding from Cancer Research UK (FC001317 and FC001175), the UK Medical Research Council (FC001317 and FC001175), and the Wellcome Trust (FC001317 and FC001175).

## DATA AND RESOURCE AVAILABILITY

All relevant data and details of resources can be found within the article and its supplementary information.

## Notes

### Competing Interest Statement

The authors have declared no competing interest.

### Summary of Updates

Additional results have been added to Figure 1 and 2, a new Figure 4 has been generated, and Supplemental files have been updated.

