## Supplementary material for "Sex-dependent plasticity of adult neural tissue in response to damage"

**SUPPLEMENTARY INFORMATION**

### Supplementary figures and legends

**Fig. S1.**

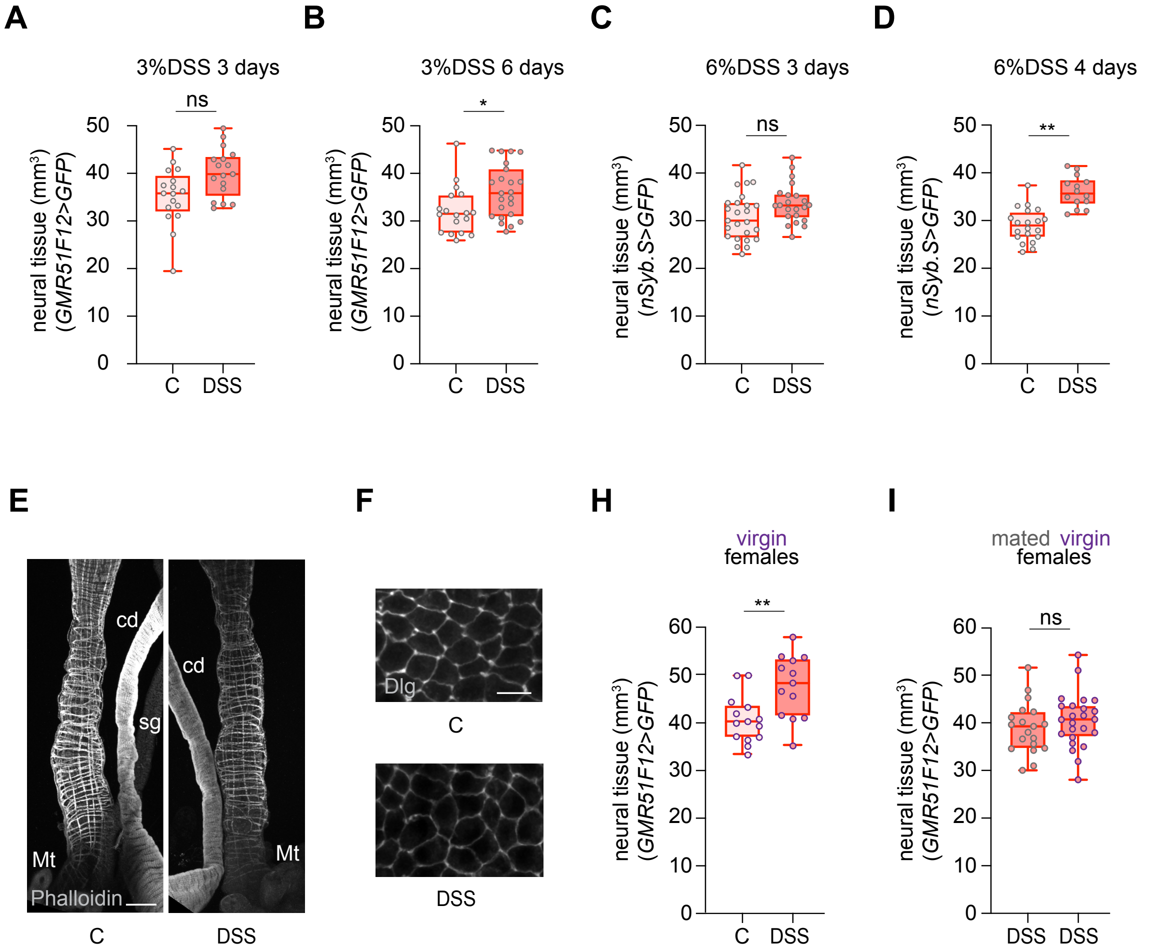


**Fig. S1. Optimization of the DSS feeding protocol resulting in neural plasticity, assessment of gut tissue integrity, and neural plasticity response in virgin flies.** A-D) Quantification of neural tissue under different DSS dosages and feeding times. A) 1 out of 4 replicates of the feeding protocol 3-day 3% DSS showed a trend towards an increase in neural tissue in the DSS-fed condition close to significance (p=0,0641). B) 1 out of 2 replicates for the feeding protocol 6 -day 3% DSS showed an increase in neural tissue, which was barely significant (p=0,0412). C) A replicate of the feeding protocol 3-day 6% DSS showed a trend towards an increase in neural tissue in the DSS-fed condition close to significance (p=0,0628). D) 4 out of 4 replicates of feeding protocol 4-day 6% DSS showed a consistent and statistically significant increase in neural tissue in the DSS-fed condition (p=0,0092 for the representative replicate shown). This is the DSS feeding protocol used throughout this study. E, F) Confocal Z-projections of the anterior midgut area, where neural tissue quantification analysis takes place, show the integrity of digestive tube tissues in the C and DSS-fed condition of 4-day 6%DSS. F) Staining with Phalloidin (grey) to visualize muscle fibers. To minimize tissue manipulation during dissection, in addition to the digestive tube, the crop duct (cd), salivary gland (sg), and Malpighian tubules (Mt) are present in the image. Scale bar 50 μm. E) Visualization of the outline of the gut epithelial cells (green, anti-Discs large (Dlg)). Scale bar 10 μm. G) Quantification of the neural tissue in virgin flies in response to C or DSS feeding. H) Quantification of neural tissue in mated and virgin flies, both DSS-fed. Statistical significance is indicated as follows: * $p < 0.05$, ** $p < 0.01$, *** $p < 0.001$, ns=not significant.

**Fig. S2.**

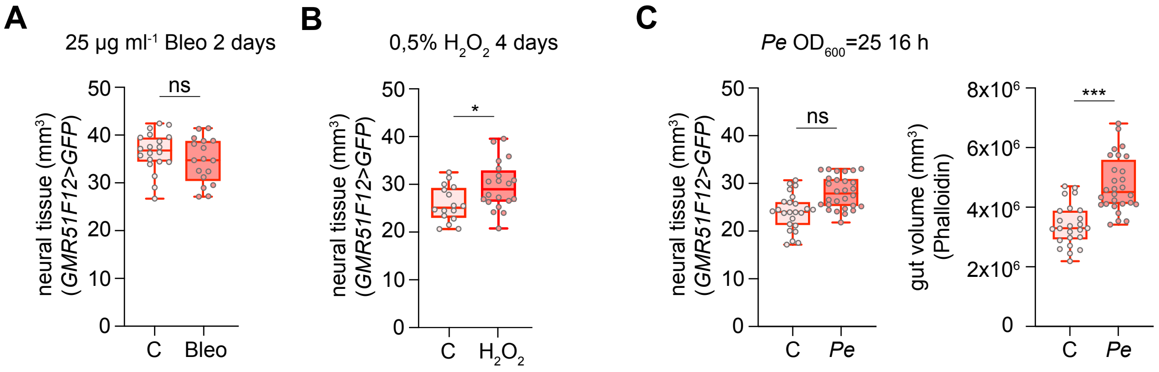


**Fig. S2. Dosage ranges and feeding times tested for Bleomycin, $H_2O_2$, and *Pseudomonas entomophila (Pe)*.** A) Quantification of the neural tissue in response to C or 2-day 25 µg $ml^{-1}$ Bleomycin (Bleo) feeding protocol. Two biological replicates of this experiment failed to show a neural increase (p=0,2275 for the replicate shown). B) Quantification of the neural tissue in response to C or 4-day 0,5% $H_2O_2$ feeding protocol. Two biological replicates of this experiment showed neural increase but with lower p value (p value=0,0357 ,for the replicate shown) than 1% (Fig. 1E, p value=0,0047). C) Quantification of neural tissue after C or *Pe* feeding $OD_{600}$=25 for 16 h. The difference between C and *Pe* is not significant (p=0,2344) due to the large increase in gut volume (p=0,0005). Statistical significance is indicated as follows: $^{*}p < 0.05$, $^{**}p < 0.01$, $^{***}p < 0.001$, ns=not significant.

**Fig. S3.**

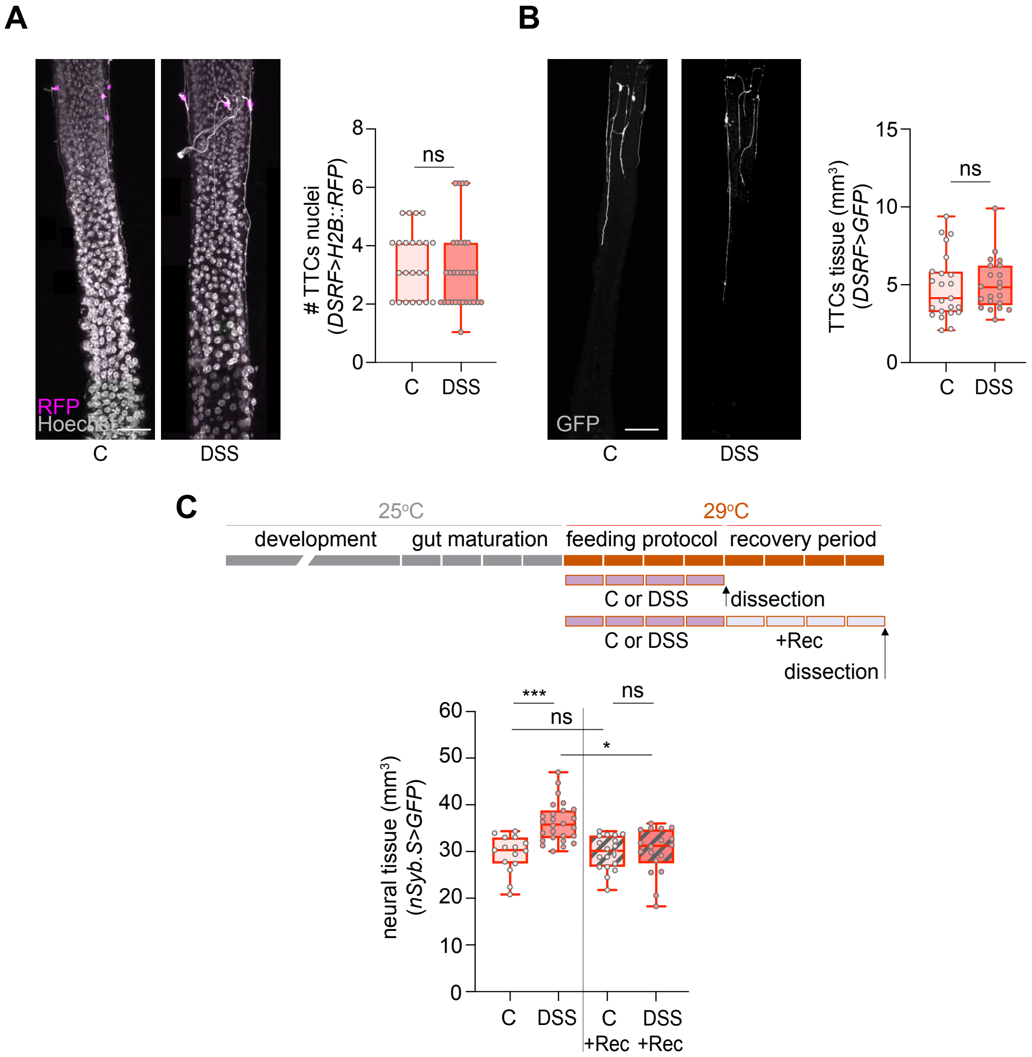


**Fig. S3. Analysis of terminal tracheal cells (TTCs) plasticity and validation of neural plasticity with the *nSyb.S-GAL4* driver.** A) Confocal Z-projections of the midgut area used to do neural tissue quantification analysis, stained with Hoechst (grey, which also accumulates in some tracheal branches) and showing the TTCs nuclei (magenta, anti-DsRed). Quantification of TTCs nuclei in C and DSS-fed females. B) Confocal Z-projections of TTCs and their branches (grey, anti-GFP) in C versus DSS-fed females. Quantification of TTCs tissue in C and DSS-fed females. C) Schematic outlining the feeding protocol, as well as the feeding protocol followed by the recovery period. Quantification of neural tissue after the feeding protocol and after the recovery period (Rec) in C and DSS-fed females using the *nSyb.S-GAL4* driver. Scale bar 50 μm. Statistical significance is indicated as follows: * $p < 0.05$, ** $p < 0.01$, *** $p < 0.001$, ns=not significant.

**Fig. S4.**

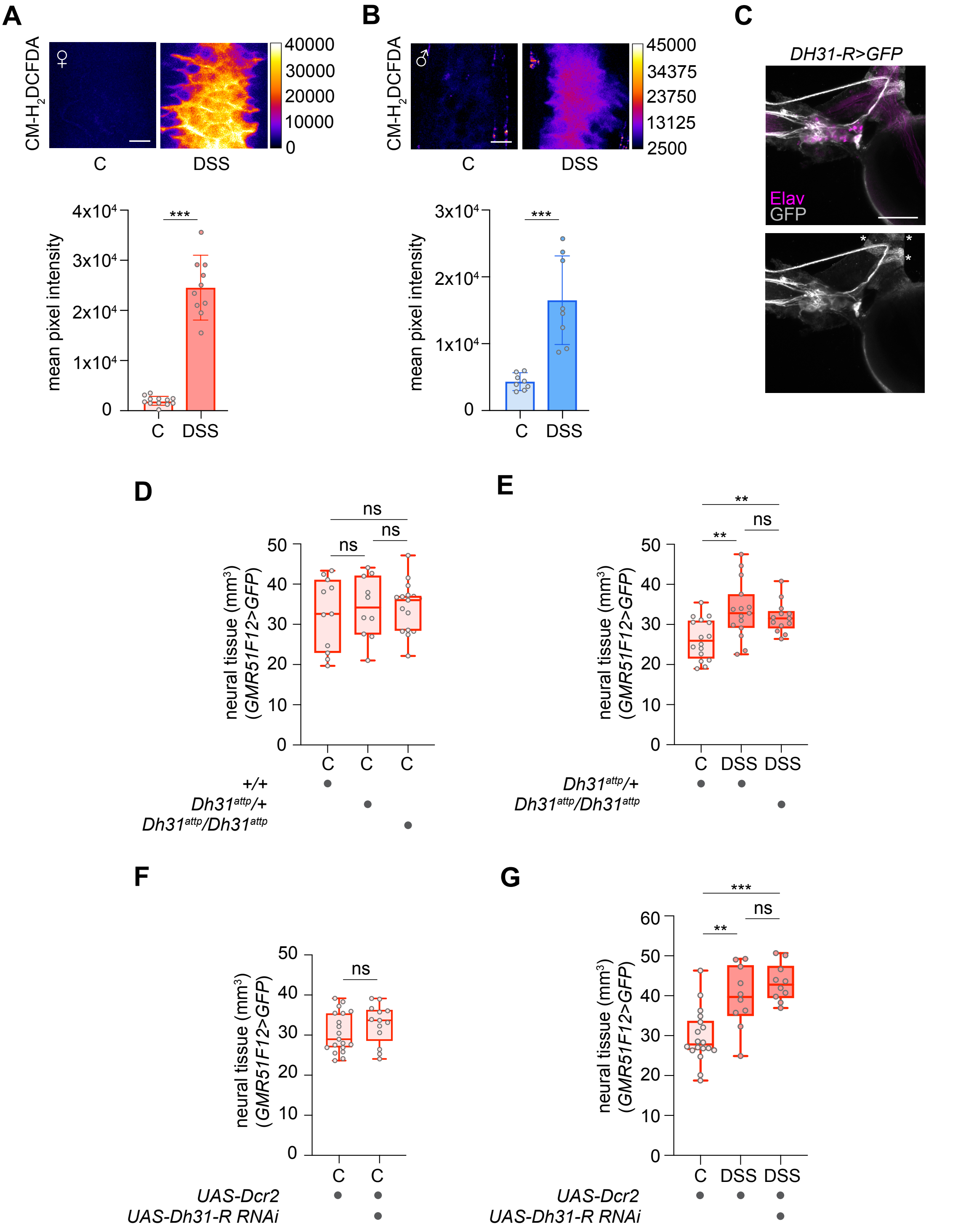


**Fig. S4. ROS-induced neural plasticity occurs independently of the Dh31/Dh31-R pathway.** A-B) DSS-dependent ROS detection in female (A) and male (B) guts. Confocal Z-projections at the level of the gut epithelium stained with CM-$H_2$DCFDA. Scale bar 20

μm. Quantification of mean pixel intensity of CM-$H_2$DCFDA signal (Fire lut) in C and DSS-fed animals. C) Confocal Z-projections showing the expression of *Dh31-R* (grey, GFP) in a subset of neuronal cell bodies (magenta, anti-Elav) in the HCG. GFP signal outlines cell bodies as well as neural projections. As reported in Medina et al. 2025, Dh31-R is also expressed in muscle cells (asterisks). Scale bar 50 μm. D) Quantification of neural tissue in wild-type, *Dh31* heterozygous and homozygous mutants in control feeding conditions. E) Quantification of neural tissue in control and DSS-fed *Dh31* heterozygous flies, to compare with the levels of neural tissue in DSS-fed *Dh31* homozygous mutant flies. F) Quantification of neural tissue in control (*Gal4/drc2* control) and neural *Dh31-R* knock-down flies in C feeding conditions. G) Quantification of neural tissue in C and DSS-fed control flies (*Gal4/drc2* control), to compare with the levels of neural tissue in DSS-fed neural *Dh31-R* knock-down flies. Statistical significance is indicated as follows: * $p < 0.05$, ** $p < 0.01$, *** $p < 0.001$, ns=not significant.

**Fig. S5.**

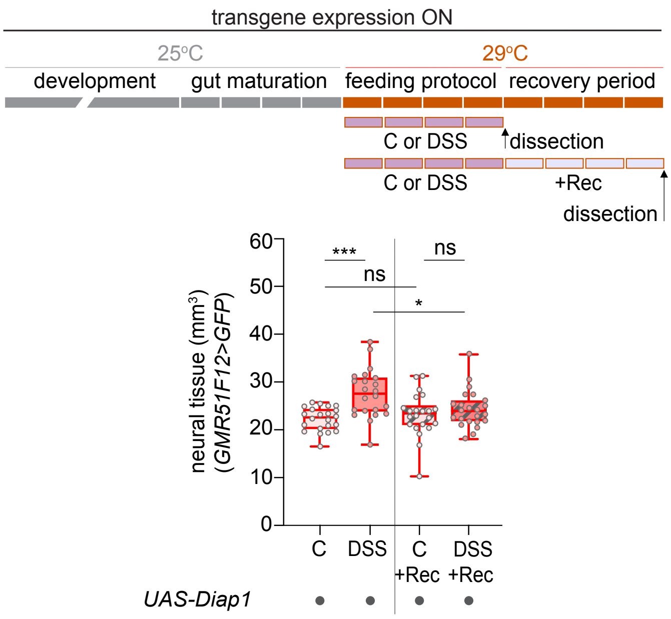


**Fig. S5. Overexpression of Diap1 does not prevent the reversal of neural growth during gut recovery.** Schematic showing transgene expression in neurons in relation to the feeding protocol and recovery period. Quantification of neural tissue after the feeding protocol and after the recovery period (Rec) in C and DSS-fed females overexpressing *Diap1*. Statistical significance is indicated as follows: *$p < 0.05$, **$p < 0.01$, ***$p < 0.001$,* ns=not significant.

__**List of genotypes**__

**Fig. 1A-E**

*w; UAS-Flybow.1.1B/UAS-Flybow.1.1B; GMR51F12-GAL4 /TM6B*

**Fig. 2A**

*w; UAS-Flybow.1.1B/UAS-Flybow.1.1B; GMR51F12-GAL4 /TM6B*

**Fig. 2B**

w; *esg-GAL4, UAS-mCD8::GFP/CyO; UAS-H2B::RFP/tubP-GAL80*$^{ts}$

**Fig. 2C-D**

*w; Cdk1*$^{null}$*/CyO; UAS-Flybow.1.1B/UAS-Flybow.1.1B, GMR51F12-GAL4*

*w; Cdk1*$^{null}$*/Cdk*$^{1E1–24}$*; UAS-Flybow.1.1B/UAS-Flybow.1.1B, GMR51F12-GAL4*

**Fig. 2E**

*w; UAS-Flybow.1.1B/UAS-Flybow.1.1B; GMR51F12-GAL4 /TM6B*

**Fig. 3A**

*w; UAS-Flybow.1.1B/UAS-Flybow.1.1B; GMR51F12-GAL4 /TM6B*

**Fig. 3B, C**

*w; UAS-Flybow.1.1B/CyO*; *traF-GAL4/UAS-Flybow.1.1B*

**Fig. 3D**

*w; UAS-Flybow.1.1B/UAS-Dcr-2; UAS-Flybow.1.1B, GMR51F12-GAL4/TM6B*

*w; UAS-Flybow.1.1B/UAS-Dcr-2; UAS-Flybow.1.1B, GMR51F12-GAL4/UAS-traRNAi*

**Fig. 3E**

*w; UAS-Flybow.1.1B/+; UAS-Flybow.1.1B, GMR51F12-GAL4/+*

*w; UAS-Flybow.1.1B/+; UAS-Flybow.1.1B, GMR51F12-GAL4/UAS-Sxl*

**Fig. 4A-B**

*w; UAS-Flybow.1.1B/UAS-Flybow.1.1B; GMR51F12-GAL4 /TM6B*

**Fig. 4C-D**

*w; UAS-Flybow.1.1B/UAS-Dcr-2; GMR51F12-GAL4/+*

*w; UAS-Flybow.1.1B/UAS-Dcr-2; UAS-traRNAi/TM6B*

*w; UAS-Flybow.1.1B/UAS-Dcr-2; GMR51F12-GAL4/ UAS-traRNAi*

**Supplementary Fig. 1A-D, G-H**

*w; UAS-Flybow.1.1B/UAS-Flybow.1.1B; GMR51F12-GAL4 /TM6B*

**Supplementary Fig. 1E-F**

Canton-S

**Supplementary Fig. 2**

*w; UAS-Flybow.1.1B/UAS-Flybow.1.1B; GMR51F12-GAL4 /TM6B*

**Supplementary Fig. 3A-B**

*w; UAS-mCD8::GFP/DSRF.Term-GAL4; UAS-H2B::RFP/TM6B*

**Supplementary Fig. 3C**

*w; UAS-Flybow.1.1B/UAS-Flybow.1.1B; nSyb-GAL4.S/TM6B*

**Supplementary Fig. 4A**

*w; Dh31-R*$^{2A\text{-}Gal4}$*/+; UAS-Flybow.1.1B/+*

**Supplementary Fig. 4B**

*w; +/+; UAS-Flybow.1.1B, GMR51F12-GAL4/ UAS-Flybow.1.1B*

*w; Dh31*$^{attP}$*/CyO; UAS-Flybow.1.1B, GMR51F12-GAL4/ UAS-Flybow.1.1B*

*w; Dh31*$^{attP}$*/ Dh31*$^{attP}$*; UAS-Flybow.1.1B, GMR51F12-GAL4/ UAS-Flybow.1.1B*

**Supplementary Fig. 4C**

*w; Dh31*$^{attP}$*/CyO; UAS-Flybow.1.1B, GMR51F12-GAL4/ UAS-Flybow.1.1B*

*w; Dh31*$^{attP}$*/ Dh31*$^{attP}$*; UAS-Flybow.1.1B, GMR51F12-GAL4/ UAS-Flybow.1.1B*

**Supplementary Fig. 4D-E**

*w; UAS-Flybow.1.1B/UAS-Dcr-2; UAS-Flybow.1.1B, GMR51F12-GAL4/TM6B*

*w; UAS-Flybow.1.1B/UAS-Dcr-2; UAS-Flybow.1.1B, GMR51F12-GAL4/UAS-Dh31-R RNAi*

**Supplementary Fig. 5**

*w; UAS-Flybow.1.1B/CyO; UAS-Flybow.1.1B, GMR51F12-GAL4/UAS-Diap1*

### Tables

**Table S1: Statistical details for Figures and Supplementary figures**

Each tab of the Excel file provides the statistical details for specific figures and supplementary figures. For each figure panel, we report the measurement performed and the experimental conditions, the number of data points (normally the number of guts analyzed unless otherwise indicated), a definition of the value measured, the statistical test used, and the predetermined comparisons performed, the resulting *p*-values, and the number of biologically independent replicates.
