## Supplementary material for "Sex-dependent plasticity of adult neural tissue in response to damage": Table S1

**Table S1**
**Statistical details for Figures and Supplementary figures**

One tab per figure

Includes: measure, experimental conditions, number of data points, definition of value, statistical test and comparisons, p value and biologically independent replicates

**Fig 1**

| PANEL | Measure and Experimental conditions | n=guts | Median | Statistics: Test | Statistics: Comparisons | p value | N=biologically independent replicates |
|---|---|---|---|---|---|---|---|
| B | **neural tissue mm3** | | | | | | |
| | C | 17 | 28,470 | ANCOVA | C vs DSS | 0,0058 | 4 |
| | DSS | 22 | 32,156 | | | | |
| C | **HCG neuronal nuclei** | | mean±s.d. | | | | |
| | C | 18 | 32,94±3,96 | unpaired t-test | C vs DSS | 0,8114 | 3 |
| | DSS | 17 | 32,65±3,29 | | | | |
| D | **neural tissue mm3** | | | | | | |
| | C | 22 | 33,241 | ANCOVA | C vs Bleo | 0,0077 | 3 |
| | Bleo | 15 | 36,008 | | | | |
| E | **neural tissue mm3** | | | | | | |
| | C | 17 | 25,399 | ANCOVA | C vs H2O2 | 0,0047 | 2 |
| | H2O2 | 23 | 30,656 | | | | |
| F | **neural tissue mm3** | | | | | | |
| | C | 19 | 30,720 | ANCOVA | C vs DSS | 0,0005 | 3 |
| | DSS | 19 | 35,592 | | DSS vs DSS+Rec | 0,0000 | |
| | C+Rec | 18 | 31,040 | | C vs C+Rec | 0,7653 | |
| | DSS+Rec | 20 | 27,548 | | C+Rec vs DSS+Rec | 0,1232 | |

Fig 2

| PANEL | Measure and Experimental conditions | n=guts | Median | Statistics: Test | Statistics: Comparisons | p value | N=biologically independent replicates |
|---|---|---|---|---|---|---|---|
| A | **PH3+ cells Anterior midgut** | | | | | | |
| | C | 24 | 0 | Mann-Whitney | C vs DSS | 0,0049 | 2 |
| | DSS | 23 | 2 | | | | |
| | **PH3+ cells All midgut** | | | | | | |
| | C | 24 | 1 | Mann-Whitney | C vs DSS | <0,0001 | 2 |
| | DSS | 23 | 21 | | | | |
| B | **% ISCs/EBs area/area total** | | **Mean±S.D.** | | | | |
| | C | 16 | 6,414±2,521 | unpaired t-test | C vs DSS | <0,0001 | 3 |
| | DSS | 18 | 16,50±7,071 | | | | |
| | **% ECs/ReDMM cells** | | | | | | |
| | C | 16 | 1,862 | Mann-Whitney | C vs DSS | 0,2879 | 3 |
| | DSS | 18 | 5,258 | | | | |
| C | **PH3+ cells Anterior midgut DSS** | | | | | | |
| | Cdk1null/+ (Cdk het) | 24 | 0 | Mann-Whitney | Cdk1 het vs Cdk1 null | 0,0018 | 2 |
| | Cdk1null/Cdk1E1-24 (Cdk null) | 21 | 0 | | | | |
| | **PH3+ cells All midgut DSS** | | | | | | |
| | Cdk1null/+ (Cdk het) | 24 | 19 | Mann-Whitney | Cdk1 het vs Cdk1 null | <0,0001 | 2 |
| | Cdk1null/Cdk1E1-24 (Cdk null) | 21 | 0 | | | | |
| D | **neural tissue mm3 DSS** | | | | | | |
| | Cdk1null/+ (Cdk het) | 23 | 24,369 | ANCOVA | Cdk1 het vs Cdk1 null | 0,8219 | 2 |
| | Cdk1null/Cdk1E1-24 (Cdk null) | 26 | 23,923 | | | | |
| E | **neural tissue mm3** | | | | | | |
| | DSS | 21 | 28,714 | ANCOVA | DSS vs DSS+NAC | 0,0034 | 3 |
| | DSS+NAC | 20 | 25,838 | | DSS+NAC vs C+NAC | 0,9687 | |
| | C+NAC | 18 | 26,006 | | | | |

**Fig 3**

| PANEL | Measure and Experimental conditions | n=guts | Median | Statistics | | p value | N=biologically independent replicates |
|---|---|---|---|---|---|---|---|
| | | | | Test | Comparisons | | |
| A | **neural tissue mm3** | | | | | | |
| | C | 24 | 27,790 | ANCOVA | C vs DSS | 0,8362 | 3 |
| | DSS | 24 | 26,950 | | | | |
| D | **neural tissue mm3** | | | | | | |
| | females C | 21 | 22,144 | ANCOVA | females C vs DSS | 0,0021 | 2 |
| | females DSS | 23 | 24,173 | | females w/ masculinized neurons C vs DSS | 0,2287 | |
| | females w/ masculinized neurons C | 24 | 21,574 | | females C vs females w/ masculinized neurons C | 0,9796 | |
| | females w/ masculinized neurons DSS | 19 | 20,042 | | females DSS vs females w/ masculinized neurons DSS | 0,0002 | |
| E | **neural tissue mm3** | | | | | | |
| | males C | 25 | 25,381 | ANCOVA | males C vs DSS | 0,9149 | 2 |
| | males DSS | 27 | 25,907 | | males w/ feminized neurons C vs DSS | 1 | |
| | males w/ feminized neurons C | 20 | 21,264 | | males C vs males w/ feminized neurons C | 0,0012 | |
| | males w/ feminized neurons DSS | 23 | 22,853 | | males DSS vs males w/ feminized neurons DSS | 0,0005 | |

**Fig 4**

| PANEL | Measure and Experimental conditions | Median | Additional Information<br>see Material and Methods<br>n=number of quadrants quantified | total flies per condition | | Test | Statistics<br>Comparisons | p value | | | N=biologically independent replicates |
|---|---|---|---|---|---|---|---|---|---|---|---|
| A | **fecal spots per quadrant/total flies** | | | | | | | | | | |
| | C | 0,711 | 8 | 38 | slipt in 2 cages | | C vs DSS | 0,0002 | | | |
| | DSS | 3,958 | 8 | 47 | slipt in 2 cages | Mann-Whitney | | | | | 2 |
| | C+Rec (sucrose) | 0,103 | 8 | 30 | slipt in 2 cages | | C+Rec vs DSS+Rec | 0,0002 | | | |
| | DSS+Rec (sucrose) | 0,818 | 8 | 22 | slipt in 2 cages | | | | | | |
| B | **fecal spots per quadrant/total flies** | | | | | | | | | | |
| | C | 0,566 | 8 | 48 | slipt in 2 cages | | C vs DSS | <0,0001 | | | |
| | DSS | 3,966 | 16 | 90 | slipt in 4 cages | Mann-Whitney | | | | | 2 |
| | C+Rec (food) | 0,333 | 8 | 47 | slipt in 2 cages | | C+Rec vs DSS+Rec | 0,1878 | | | |
| | DSS+Rec (food) | 0,731 | 16 | 77 | slipt in 4 cages | | | | | | |
| C | **fecal spots per quadrant/total flies** | | | | | | | | | | |
| | females Gal4/drc2 control C | 0,477 | 8 | 52 | slipt in 2 cages | | Gal4/drc2 control C vs DSS | <0,0001 | | | |
| | females Gal4/drc2 control DSS | 1,159 | 12 | 71 | split in 3 cages | | | | | | |
| | females traRNAi/dcr2 control C | 0,944 | 8 | 48 | slipt in 2 cages | Mann-Whitney | traRNAi/dcr2 control C vs DSS | <0,0001 | | | 3 |
| | females traRNAi/dcr2 control DSS | 2,083 | 12 | 69 | split in 3 cages | | | | | | |
| | females w/ masculinized neurons C | 1,978 | 8 | 52 | slipt in 2 cages | | females w/ masculinized neurons C vs DSS | 0,1013 | | | |
| | females w/ masculinized neurons DSS | 2,403 | 12 | 77 | split in 3 cages | | | | | | |
| D | **survival probability curves** | **Number of flies** | | | | | | **HR** | **log-rank test p value** | **p value adjusted** | |
| | females Gal4/drc2 control C | 100 | | | | | females Gal4/drc2 control C vs DSS | 6,1904 | <0,001 | <0,001 | |
| | females Gal4/drc2 control DSS | 135 | | | | | females traRNAi/dcr2 control C vs DSS | 3,8980 | <0,001 | <0,001 | |
| | females traRNAi/dcr2 control C | 100 | | | | | females w/ masculinized neurons C vs DSS | 27,0529 | <0,001 | <0,001 | |
| | females traRNAi/dcr2 control DSS | 112 | | | | | | | | | 3 |
| | females w/ masculinized neurons C | 90 | | | | log-rank test with Holm-Bonferroni correction | females Gal4/drc2 control DSS vs females w/ masculinized neurons DSS | 4,7001 | <0,001 | <0,001 | |
| | females w/ masculinized neurons DSS | 112 | | | | | females traRNAi/dcr2 control DSS vs females w/ masculinized neurons DSS | 5,3879 | <0,001 | <0,001 | |
| | | | | | | | females Gal4/drc2 control C vs females w/ masculinized neurons C | 1,4169 | 0,0900 | 0,6590 | |
| | | | | | | | females traRNAi/dcr2 control C vs females w/ masculinized neurons C | 1,2698 | 0,3000 | 0,9990 | |

Fig S1

| PANEL | Measure and Experimental conditions | n=guts | Median | Statistics | | p value | N=biologically independent replicates |
|---|---|---|---|---|---|---|---|
| | | | | Test | Comparisons | | |
| A | **3%DSS 3days** | | | | | | |
| | **neural tissue mm3** | | | | | | |
| | C | 17 | 35,820 | ANCOVA | C vs DSS | 0,0641 | 4 |
| | DSS | 17 | 39,859 | | | | |
| B | **3%DSS 6days** | | | | | | |
| | **neural tissue mm3** | | | | | | |
| | C | 18 | 31,582 | ANCOVA | C vs DSS | 0,0412 | 1 |
| | DSS | 23 | 35,905 | | | | |
| C | **6%DSS 3days** | | | | | | |
| | **neural tissue mm3** | | | | | | |
| | C | 25 | 29,912 | ANCOVA | C vs DSS | 0,0628 | 1 |
| | DSS | 22 | 33,054 | | | | |
| D | **6%DSS 4days** | | | | | | |
| | **neural tissue mm3** | | | | | | |
| | C | 20 | 28,797 | ANCOVA | C vs DSS | 0,0092 | 4 |
| | DSS | 14 | 35,481 | | | | |
| G | **virgin females** | | | | | | |
| | **neural tissue mm3** | | | | | | |
| | C | 14 | 40,105 | ANCOVA | C vs DSS | 0,0040 | 2 |
| | DSS | 13 | 48,151 | | | | |
| H | **neural tissue mm3 DSS** | | | | | | |
| | mated females | 19 | 39,369 | ANCOVA | mated vs virgin females | 0,4386 | 2 |
| | virgin females | 24 | 40,862 | | | | |

Fig S2

| PANEL | Measure and Experimental conditions | n=guts | Median | Statistics | | p value | N=biologically independent replicates |
|---|---|---|---|---|---|---|---|
| | | | | Test | Comparisons | | |
| A | **neural tissue mm3** | | | | | | |
| | C | 20 | 36,795 | ANCOVA | C vs Bleo | 0,2275 | 2 |
| | Bleo | 17 | 34,792 | | | | |
| B | **neural tissue mm3** | | | | | | |
| | C | 16 | 25,100 | ANCOVA | C vs $H_2O_2$ | 0,0357 | 2 |
| | $H_2O_2$ | 20 | 28,902 | | | | |
| C | **neural tissue mm3** | | | | | | |
| | C | 24 | 24,079 | ANCOVA | C vs *Pe* | 0,2344 | 1 |
| | *Pe* | 28 | 27,948 | | | | |
| | | | | gut volume covariate p value | | 0,0005 | |

Fig S3

| PANEL | Measure and Experimental conditions | n=guts | Median | Statistics | | p value | N=biologically independent replicates |
|---|---|---|---|---|---|---|---|
| | | | | Test | Comparisons | | |
| A | **TTCs nuclei** | | | | | | |
| | C | 23 | 3,000 | Mann-Whitney | C vs DSS | 0,2825 | 2 |
| | DSS | 31 | 3,000 | | | | |
| B | **TTCs tissue mm3** | | | | | | |
| | C | 23 | 4,139 | ANCOVA | C vs DSS | 0,6928 | 2 |
| | DSS | 21 | 4,849 | | | | |
| C | **neural tissue mm3** | | | | | | |
| | C | 15 | 30,314 | ANCOVA | C vs DSS | 0,0000 | 3 |
| | DSS | 26 | 35,761 | | DSS vs DSS+Rec | 0,0128 | |
| | C+Rec | 18 | 30,154 | | C vs C+Rec | 0,2089 | |
| | DSS+Rec | 18 | 31,233 | | C+Rec vs DSS+Rec | 0,1053 | |

Fig S4

| PANEL | Measure and Experimental conditions | n=guts | Mean±S.D. | Statistics | | p value | N=biologically independent replicates |
|---|---|---|---|---|---|---|---|
| | | | | Test | Comparisons | | |
| A | **mean pixel intensity ROS signal females** | | | | | | |
| | C | 12 | 1995±890,4 | unpaired t-test | C vs DSS | <0,0001 | 2 |
| | DSS | 8 | 24546±6465 | | | | |
| B | **mean pixel intensity ROS signal males** | | | | | | |
| | C | 6 | 4325±1337 | unpaired t-test | C vs DSS | 0,0009 | 2 |
| | DSS | 8 | 16505±6621 | | | | |
| | | | **Median** | | | | |
| D | **neural tissue mm3 C** | | | | | | |
| | wild type | 11 | 32,618 | | wild type vs Dh31attp/+ | 0,4682 | |
| | Dh31attp/+ | 10 | 34,179 | ANCOVA | wild type vs Dh31attp/Dh31attp | 0,1005 | 1 |
| | Dh31attp/Dh31attp | 17 | 36,032 | | Dh31attp/+ vs Dh31attp/Dh31attp | 0,8432 | |
| E | **neural tissue mm3** | | | | | | |
| | Dh31attp/+ C | 16 | 25,945 | | Dh31attp/+ C vs DSS | 0,0043 | |
| | Dh31attp/+ DSS | 15 | 32,799 | ANCOVA | Dh31attp/+ DSS vs Dh31attp/Dh31attp DSS | 0,2498 | 2 |
| | Dh31attp/Dh31attp DSS | 13 | 31,503 | | Dh31attp/+ C vs Dh31attp/Dh31attp DSS | 0,0051 | |
| F | **neural tissue mm3 C** | | | | | | |
| | UAS-Dcr2 control | 19 | 28,986 | ANCOVA | UAS-Dcr2 control vs Dh31 R RNAi expression | 0,2254 | 1 |
| | Dh31 R RNAi expression | 13 | 33,688 | | | | |
| G | **neural tissue mm3** | | | | | | |
| | UAS-Dcr2 control C | 18 | 27,766 | | UAS-Dcr2 control C vs DSS | 0,0024 | |
| | UAS-Dcr2 control DSS | 10 | 39,689 | ANCOVA | UAS-Dcr2 control DSS vs Dh31 R RNAi expression DSS | 0,6689 | 2 |
| | Dh31 R RNAi expression DSS | 10 | 42,71 | | Dh31attp/+ C vs Dh31attp/Dh31attp DSS | 0 | |

Fig S5

| PANEL | Measure and Experimental conditions | n=guts | Median | Statistics: Test | Statistics: Comparisons | p value | N=biologically independent replicates |
|---|---|---|---|---|---|---|---|
| | **neural tissue mm3** | | | | | | |
| | C | 22 | 22,594 | ANCOVA | C vs DSS | 0,0002 | 2 |
| | DSS | 22 | 27,566 | | DSS vs DSS+Rec | 0,0216 | |
| | C+Rec | 22 | 23,334 | | C vs C+Rec | 0,5686 | |
| | DSS+Rec | 23 | 24,058 | | C+Rec vs DSS+Rec | 0,3096 | |
